# Distinct genomic and epigenomic features demarcate hypomethylated blocks in colon cancer

**DOI:** 10.1101/028803

**Authors:** Mahfuza Sharmin, Héctor Corrada Bravo, Sridhar Hannenhalli

## Abstract

**Background.** Large mega base-pair genomic regions show robust alterations in DNA methylation levels in multiple cancers, a vast majority of which are hypo-methylated in cancers. These regions are generally bounded by CpG islands, overlap with *Lamin Associated Domains* and *Large organized chromatin lysine modifications,* and are associated with stochastic variability in gene expression. Given the size and consistency of *hypo-methylated blocks (HMB)* across cancer types, their immediate causes are likely to be encoded in the genomic region near HMB boundaries, in terms of specific genomic or epigenomic signatures. However, a detailed characterization of the HMB boundaries has not been reported.

**Method.** Here, we focused on ~13k HMBs, encompassing approximately half the genome, identified in colon cancer. We analyzed a number of distinguishing features at the HMB boundaries including transcription factor (TF) binding motifs, various epigenomic marks, and chromatin structural features.

**Result.** We found that the classical promoter epigenomic mark – H3K4me3, is highly enriched at HMB boundaries, as are CTCF bound sites. HMB boundaries harbor distinct combinations of TF motifs. Our Random Forest model based on TF motifs can accurately distinguish boundaries not only from regions inside and outside HMBs, but surprisingly, from active promoters as well. Interestingly, the distinguishing TFs and their interacting proteins are involved in chromatin modification. Finally, HMB boundaries significantly coincide with the boundaries of *Topologically Associating Domains* of the chromatin.

**Conclusion.** Our analyses suggest that the overall architecture of HMBs is guided by pre-existing chromatin architecture, and are associated with aberrant activity of promoter-like sequences at the boundary.

## Background

Cells in an individual adopt hundreds of distinct phenotypes in their structure and function. This dramatic phenotypic variability through development and disease cannot be explained by genetic differences alone, but is rather encoded in the so-called epigenetic variation – varying degrees of chemical modifications of the DNA and nucleosome histones that the genomic DNA is wrapped around [1, 2], Epigenetic mechanisms are integral to gene regulation; and, their role in cellular differentiation [3], aging [4] and disease [5] are areas under active investigation. DNA methylation is one of the earliest known epigenetic modifications, for which cellular inheritance mechanisms are now well understood [6]. Although a direct relationship between locus-specific DNA methylation and gene expression is well known, a more specific involvement of DNA methylation in various diseases, in particular, cancer, is only beginning to be investigated in a comprehensive manner [5, 7, 8]. Collectively, these studies have identified specific oncogenes that are hypomethylated, and thus activated, in cancer [9], certain tumor suppressor genes that are hypermethylated, and thus inactivated [10], and additional methylation changes in cancer [7, 8].

A recent study showed well-demarcated, large regions, collectively covering half of the genome, to be differentially methylated in cancer [5]. Moreover, presence of such large cancer-specific differentially methylated regions (cDMRs) was found to be a general epigenomic signature across many cancer types [5]. The cDMRs contain important genes involved in mitotic cell cycle and matrix remodeling and were shown to exhibit extreme gene expression variability. Moreover, cDMRs are highly enriched among regions differentially methylated during stem cell reprogramming of induced pluripotent stem cells [11]. Subsequent investigations revealed that cDMRs significantly overlapped with Lamina Attachment Domains (LAD), Large organized chromatin lysine modifications (LOCK) [12] and Partially Methylated Domains (PMD) in cancer [3]. Additionally, 1kb regions flanking cDMR boundaries were shown to be enriched for DNase hypersensitive sites [13]. Nucleosomes were found to be locally enriched in hypomethylated regions in normal tissue [14], Collectively, these observations led the authors to postulate a model of cancer progression involving epigenetic instability of well-defined genomic domains [5]. However, investigations of additional genomic and epigenomic correlates of cDMRs, and ultimately the causes of cDMR formation are needed to gain a better mechanistic understanding of the role of DNA methylation in cancer, in order to harness the full potential of these earlier studies for epigenetic-based cancer diagnostics [15].

A vast majority of large cDMRs are in fact hypomethylated in cancer, i.e. less methylated in cancer tissue than the corresponding normal tissue, and such hypomethylation happens in large contiguous genomic regions called hypomethylated blocks. Here, we focused on ~13k hypomethylated blocks (HMB), encompassing approximately half the genome, previously identified in colon cancer [5]. Given the length of HMBs and their general overlap with chromatin structural features such as LADs and enrichment of DNAse hypersensitive sites at HMB boundaries, it is likely that the genome and the epigenome at HMB boundaries hold the clues to the underlying mechanisms of genome wide hypomethylation with distinct boundaries. We therefore analyzed a number of genomic and epigenomic features at the HMB boundaries including TF binding motifs, epigenomic marks, and three-dimensional chromatin structural features (Figure. 1).

**Figure 1.**
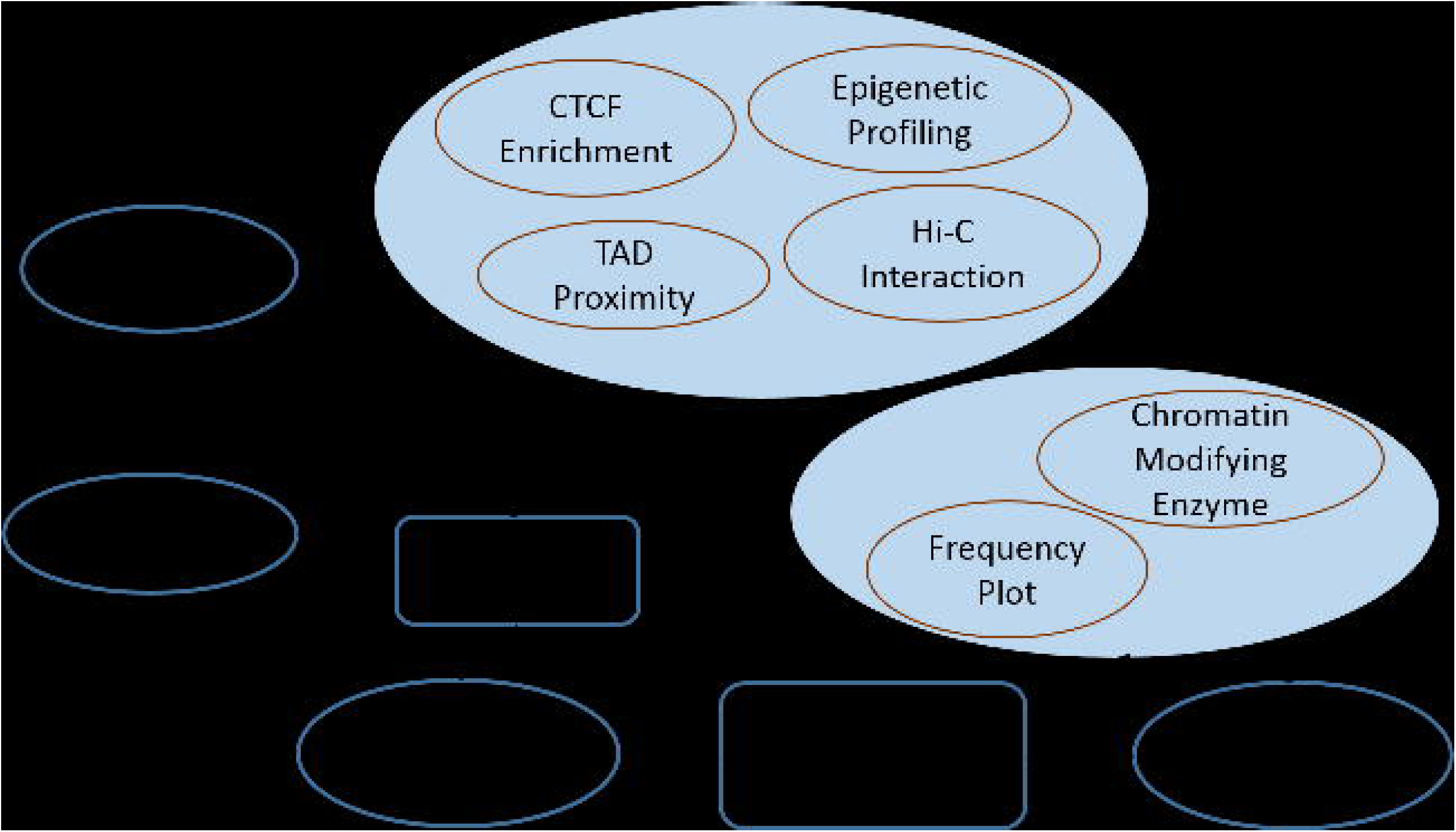
Analysis pipeline. Starting with ~13,000 HMBs, we perform a number of tests to assess the association of HMBs and HMB boundaries with Topological Associating Domains, Physical interaction within and across HMBs, profiles of various epigenetic marks, and CTCF binding. In addition, we identified TF motifs enriched at the HMB boundaries relative to various controls and assessed the ability of a random forest model to distinguish HMB boundaries from other domains based on TF binding site motifs. Finally, we assessed the spatial profile and functions of enriched TF motifs and their interacting partners.

Our analysis revealed that the classical promoter epigenomic mark – H3K4me3, is highly enriched at HMB boundary in normal colon tissue, and the boundaries that are enriched for promoter marks are also enriched for *in vivo* binding of the insulator protein CTCF in colon cancer. We also found that the HMB boundaries harbor distinct combinations of TF motifs. Our *Random Forest* machine learning model that uses TF motifs as features can distinguish boundaries not only from regions inside and outside HMBs, but surprisingly, from active promoters as well, with very high accuracy (F-measure ~ 0.98). Interestingly, the TFs that preferentially bind at HMB boundaries and their interacting partners are involved in chromatin modification. Finally, we found that HMB boundaries are associated with the boundaries of Topological Associating Domains (TADs), which form the backbone of chromatin structure [16].

Taken together, our analyses suggest that the overall architecture of HMBs is guided and restricted by pre-existing chromatin architecture, while their creation in cancer may be caused by aberrant activity of promoter-like sequences at the boundary, with a direct chromatin modification activity.

## Methods

### Hypomethylated blocks

We obtained coordinates for 13,540 reported long hypo-methylated block (HMB) in colon cancer with an average and median size of 144 kbps and 39.5 kbps, respectively [5]. We define the boundary of an HMB as its 5kb flanking regions outside the HMB plus an additional 1kb inside the HMB. The choice of 5kb for the flanking region is arbitrary and 1kb inside is included to offset a lack of precision in localizing HMB boundary (e.g., Supplementary Figure 10b of [5]).

### Random Forest based discrimination of HMB boundaries

We used Random Forest classifiers [17] to distinguish the resulting 27,080 6-kb-long HMB boundary from other genomic regions: (1) *inside HMB* - randomly selected 6kb block from inside of the HMBs, excluding HMB boundaries; (2) *outside HMB* - randomly selected 6kb regions from outside of the HMBs excluding HMB boundaries; (3) *promoter* - randomly selected 6kb promoters for protein-coding genes, including 5 kb upstream and 1 kb downstream of the transcription start site using the Ensembl annotation (www.ensembl.org, version 69). Given two sets of sequences (e.g., HMB and inside-HMB), and a set of characteristics (i.e. features) describing each sequence (e.g., putative binding sites for a set of transcription factors), the Random Forest classifier learns the combinations of features that distinguish one set of sequences from the other. When given an unforeseen sequence and its features, our Random Forest classifier can determine the set to which the sequence belongs to based on its features. The more distinguishing the features of the two sequence sets are (e.g., HMB and inside-HMB), the higher the accuracy with which our classifier can determine the set to which a new sequence belongs. To design the right control while building the Random Forest classifier, in each sequence set we selected the same numbers of regions for each pairwise classification task, while controlling for the GC content. For instance, when classifying between HMB boundaries and promoters, we selected two sets of regions that are non-overlapping and with similar GC content distribution. Finally, each set of sequences were composed of ~20k sequences.

As feature sets in the Random Forest classifiers, 931 motifs corresponding to vertebrate TFs were obtained from TRANSFAC v2011 [18]. Putative motif binding was determined in each 6kb region using the FIMO (Find individual Motif Occurrences) software [19]. Each 6 kb region was represented as a 931-dimensional feature vector where the measurement of each dimension is the count (0 or greater) of binding sites of each corresponding motif within the 6kb region. We assessed the classification accuracy using a 70%-30% split of the data into training and test sets, chosen randomly, for cross-validation in each of the pairwise classification tasks distinguishing HMB boundaries from the three sets of regions: inside HMB, outside HMB, and promoters. The classification accuracies are reported using both area under curve (AUC) of the receiver operating curve and harmonic mean of precision and recall (F-measure). As an additional robustness measure, we also performed the HMB boundary versus promoter classification task using Support Vector Machine (SVM) implemented in R statistical package (*www.r-project.org*), based on 10-fold cross-validation.

### CpG island overlap as an additional feature in the Random Forest Classifiers

CpG Islands tend to exhibit increased methylation in colon cancer. Consequently, HMBs are frequently ‘broken’ by CpG Islands [5], and thus their boundaries frequently overlap CpG islands. Therefore, motifs can be found more frequently in HMB boundaries than inside or outside HMBs simply due to the presence of CpG islands. We used the fraction of the 6kb region that overlaps any of the 28,681 CpG islands annotated in the UCSC genome browser (genome.ucsc.edu) as an additional feature in the classification task, in addition to controlling for GC content in the classification task.

### Identifying most discriminating motifs

We determined the importance of each motif in distinguishing between region types using *Mean Decrease Accuracy* obtained from the Random Forest classifier. Mean decrease accuracy of a feature measures the reduction in classification error upon including the corresponding feature in the model, and thus represents the importance of the motif in distinguishing HMB boundaries from a specific control region set; the higher the mean decrease accuracy the more important the feature is. We also determined enrichment of each motif in HMB boundaries relative to each control set (inside, outside, or promoters) using Fisher’s exact test. The motif is considered as enriched (depleted) in the HMB boundaries relative to the control when the corresponding odds ratio is greater than 2 (less than 0.5).

### Epigenetic data processing

Genome-wide profiles of six histone marks (H3K4me1, H3K4me3, and H3K9ac, H3K9me3, H3K27me3 and H3K36me3) in normal colon mucosa tissue were downloaded from the Roadmap Epigenetics Project website (www.roadmapepigenomics.org/). We calculated average signal for each histone mark (at 20bp resolution as provided by the Roadmap project) within each 6kb region in HMB, inside HMB, outside HMB, and promoter region. ChIP-Input was also obtained for normalization. To get the normalized values, we took the log ratio of methylation levels of histone marks and their corresponding ChIP-Input at the base-pair resolution.

### Chromatin interaction measurement in hypo-methylated blocks

To obtain the chromatin interaction information, we used Hi-C experimental data, which provides the spatial proximity information between pairs of different genome segments [20]. We obtained Hi-C data for human embryonic stem cell (hESC) and lung fibroblasts (hIMR90) cell lines from [16] as normalized interaction matrices with 40 kb bin size denoting the frequencies of physical contacts among pairs of genomic loci at a genome-wide scale. We mapped those 40 kb bins onto the HMBs and disregarded partially mapped blocks so HMBs smaller than 40kb were excluded from the analysis. We then measured interaction strength within each HMB as the sum of all pairwise bin interactions within the HMB divided by the number of 40 kb bins within the HMB. As a negative control, the same was done for randomly chosen non-overlapping genomic regions with same lengths as HMBs.

### Measuring Proximity to Topologically Associating Domains

We downloaded the locations of 3,029 topological associating domains (TADs) from [16] for hESC cell lines. For each boundary of the TAD we obtain the minimum distance to a HMB boundary. As a control, we selected 13k random non-overlapping blocks of same sizes as HMBs. As for real HMBs, we also obtained the minimum distance of each TAD boundary to a random block selected for control.

### Fisher’s exact test: calculating enrichment/depletion of motif in different regions and finding motif interaction with chromatin modification enzymes (CME)

The contingency table for testing enrichment/depletion of each motif is shown below.

|  | Positive | Negative |
| --- | --- | --- |
| Presence | a | c |
| Absence | b | d |

*a (*respectively *b)* denotes the number of positive examples in which a motif is present (respectively absent). Similarly, *c* (respectively *d*) denotes the number of negative examples in which a motif is present (respectively absent).

The contingency table for testing interaction with CME is shown below.

|  | Selected Motifs | Other Motifs |
| --- | --- | --- |
| Interact with CME | a | c |
| Do not interact with a CME | b | d |

*a* (respectively *b*) denotes the number of selected motifs that themselves are CMEs or do not interact with a CME (respectively all others). Similarly, *c* (respectively *d*) corresponds to the control for testing CMC interaction using all the other motifs that themselves are not CMEs.

## Results

**Overview.** Our objective is to characterize genetic and epigenetic features that demarcate Hypomethylated blocks in cancer, in order to gain insights into the mechanism and functional implications of these genomic blocks. Our findings are organized as follows: First, we determined and examined epigenomic marks that are enriched at HMB boundaries. Second, we analyzed genomic properties, namely, putative binding sites for all vertebrate transcription factors at HMB boundaries. Third, we showed that many of the motifs enriched at HMB boundaries exhibit specific positional distributions aligned with the HMB boundary. Fourth, we investigated specific transcription factor motifs enriched at HMB boundaries and their links to chromatin modifying enzymes (CMEs), in order to understand the mechanistic link between transcription factor binding and chromatin structure. Fifth, we furthered examined the link between genetic/epigenetic properties of the HMB boundaries and CMEs by analyzing the association between HMB boundaries and topologically associating domains (TAD) boundaries, which define the structural backbone of the chromatin. Finally, we examined at HMB boundaries, the putative sites for CTCF which acts both as mediator of chromatin loop formation as well as an insulator that restricts the spread of chromatin marks.

### Boundaries of hypo-methylated blocks are enriched for promoter-associated histone mark H3K4me3

Previous studies have shown cross-talk between DNA methylation and various histone modifications [21]. Given that HMBs exhibit relatively sharp demarcation of their boundaries [5], we investigated the patterns of various histone marks in normal colon tissue in the vicinity of HMB boundaries. We summarized the signal strength of six histone marks in 20 kbp flanking the HMB boundaries (see Methods) from human colon tissue data downloaded from the Epigenetic Roadmap Website (www.roadmapepigenomics.org). Histone marks H3K4me3 and H3K9ac, known to be associated with active promoters, showed a distinct peak immediately outside the HMBs (Figure 2). Patterns for other histone marks (H3K4me1, H3K9me3, H3K27me3 and H3K36me3) did not show noticeable trends at HMB boundaries (Supplementary Figure 1a-d).

**Figure 2.**
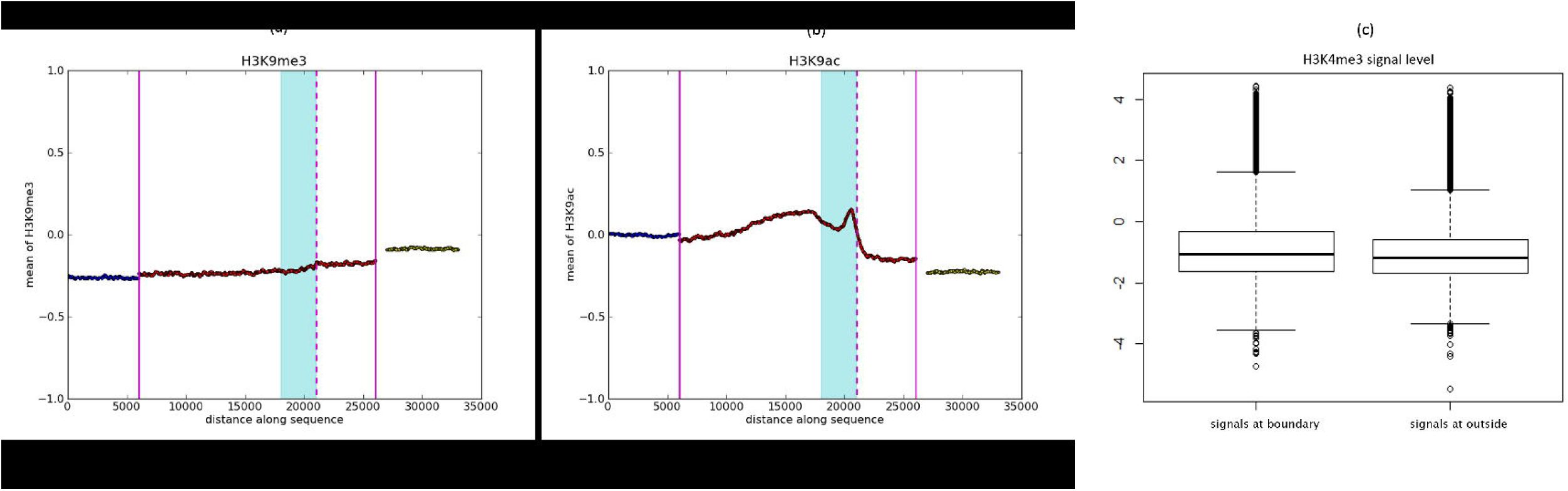
Histone modifications enriched near HMB boundaries. Mean normalized ChIP signal for *(a) H3K4me3* and *(b) H3K9ac* as a function of genomic distance to HMB boundary. The dotted vertical line (pink) depicts the precise location where the HMB starts while the shaded (cyan) region is the 3kb HMB boundary region as defined in this paper. The solid vertical lines (pink) indicate inside (right) and outside (left) of HMBs. (c) Distribution of normalized H3K4me3 signal in HMB boundary regions and outside HMBs.

Given the enrichment for promoter histone marks at the HMB boundaries, we considered the possibility that the HMB boundaries coincide with or are near gene promoters. We excluded the HMB boundaries (5kb outside the HMB and 1 kb inside the HMB) that overlapped with the transcription start site of any gene or pseudogene (including non-coding genes), based on Gencode annotation [22], and repeated the analysis of histone mark pattern. The remaining boundaries still showed a significant, but smaller than previously mentioned peak, at the HMB boundary. For instance, as shown in Figure 2c, H3K4me3 signal strength in 3kb *outside HMBs* was lower than that in the regions immediately outside HMBs. The mean of normalized signals (see Methods) at the HMB boundaries was -0.82, while at random 3kb regions outside of HMBs the mean signal was -1.01 (Wilcoxon test p-value = 7.08e-42). This suggests that the observed enrichment of histone modification at HMB boundaries is not entirely due to annotated promoters for genes or pseudogenes.

### HMB boundaries harbor distinguishing TF binding motifs

Given the enrichment for promoter-like histone marks near HMB boundaries, we assessed whether HMB boundaries are distinct from non-boundary regions as well as other known promoters in terms of their TF binding motifs. For this purpose, in addition to the HMB boundary regions we defined three sets of regions of 6kb length (see Methods): (1) *Inside:* regions within HMBs, (2) *Outside:* regions between HMBs, and (3) *Promoters.* All regions were non-overlapping and in each pairwise comparison task, the GC content was similar in the two sets of regions (See Methods). For each 6kb region we constructed a 932-dimensional feature set quantifying the fraction of CpG Island overlaps and the number of motif matches for each of the 931 vertebrate TF motifs from TRANSFAC, v2011 [18], using FIMO [19] as the motif search tool. We then applied Random Forest (RF) classifiers on the feature set to distinguish HMB boundaries from the other genomic region sets under study. We trained the RF using 70% of the data and noted the classification accuracy on the remaining 30% of the data. The classification performances are shown in Table 1. Surprisingly, HMB boundaries can be distinguished from even other promoters with very high accuracy (F-measure ~ 0.978); Figure 3 shows the ROC curve corresponding to classification between HMB boundaries and promoters (ROC curves for the rest of the classification tasks are presented in Supplementary Figure 2). We were able to recapitulate the RF results of HMB boundary versus promoter classification accuracy using Support Vector Machine (SVM) (F-measure ~0.97) – SVM is a classic tool for learning the combination of features of set of sequences that distinguishes the set from the control set. This suggests that the motif composition at HMB boundaries is distinct from those in promoter regions. We also obtained high discriminative performance when distinguishing HMB boundaries from regions inside HMBs (F-measure ~0.90).

**Table 1.** Performance of Random Forest classifier for HMB boundaries relative to other genomic regions. ‘Inside’ and ‘outside’ refer to regions inside or outside HMBs, respectively. These regions were selected to match the length and CG content of HMB boundaries (see Methods). The last row contains the results of a Support Vector Machine classifier that was used to replicate the Random Forest result on the HMB boundary vs. Promoter region classification. In all cases, 70% of the data was used as training, and 30% was used for testing. Sensitivity, Specificity and F-measure were noted as the optimal F-measure.

|  | Sensitivity | Specificity | F-measure | AUC | Size of Data Set |
| --- | --- | --- | --- | --- | --- |
| Boundary vs. Inside | 0.90 | 0.89 | 0.90 | 0.96 | 41,425 |
| Boundary vs. Outside | 0.84 | 0.81 | 0.83 | 0.91 | 41,430 |
| Boundary vs. Promoter | 0.98 | 0.97 | 0.98 | 0.99 | 31,051 |
| Boundary vs. Promoter (SVM) | 0.97 | 0.97 | 0.97 | 0.99 | 31,051 |

**Figure 3.**
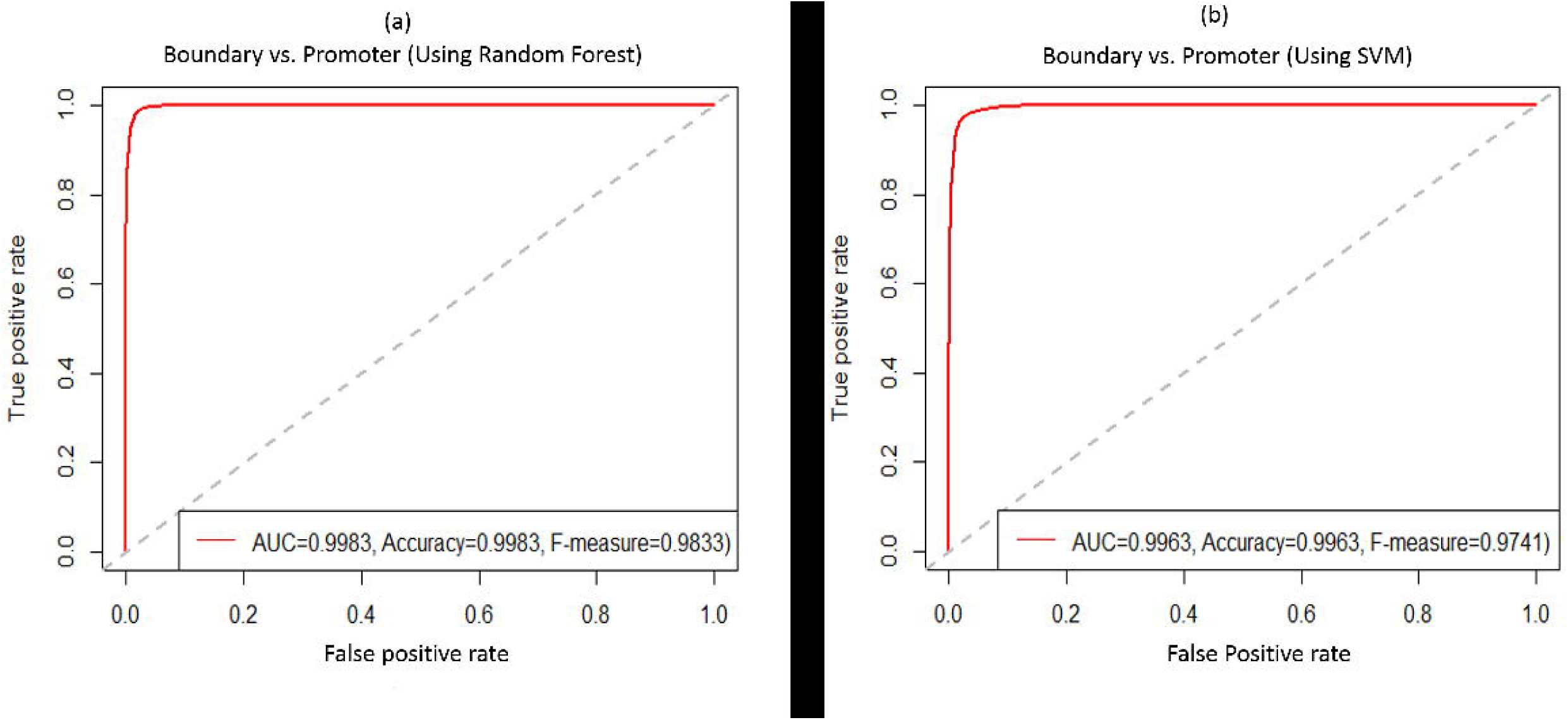
ROC curves for classifiers distinguishing HMB boundaries and promoters based on TF binding site motifs. (a) Using Random Forest classifier, (b) Using Support Vector Machine classifier. Each ROC curve is based on predictions on a held-aside set of genomic regions (see Methods).

### Positional distribution of discriminating motifs

Next we assessed whether TF motifs that distinguish HMBs exhibit a positional bias relative to the HMB boundaries. To prioritize the motifs we used the Mean Decrease Accuracy as the measure of a motif’s relevance to a specific discrimination task (see Methods). Supplementary Table 4 lists the top 20 most discriminating motifs in the classification of HMB boundaries against inside-HMB, outside-HMB, and promoters. Also, we only selected 46 motifs that were enriched above a threshold in the boundary (see Methods). For each of the 46 motifs, we plotted the frequency of the motif in 100 bps windows within the 6 kb HMB boundary regions, averaged over all HMB boundaries. Figure 4 shows the positional profile for the two most discriminating transcription factors ZFX (TransFac id M01593) and SP1 (TransFac id M00196) as an illustration; the profiles of all other motifs are included in Supplementary Figure 3. We next estimated for each motif the positional bias of binding sites within HMB boundaries by taking the most extreme (high or low) frequency of binding motifs among all 100 bp windows. The extreme frequencies of binding motifs were normalized and converted to Z-scores across all 100 bp windows in the 6kb regions. Z-score provides a standardized measurement of deviation from the mean frequency of binding motifs across the 46 motifs. We found that the majority of extreme frequency was located near the HMB boundaries: within 6k block the median location is 5574 from the outside of the boundary with a standard deviation of 892. Z-scores for all motifs ranged from 2.35 to 5.94 with a mean of 3.48 (See Supplementary Figure 3 for all positional profiles, the corresponding Z-score for both boundary and promoter), suggesting that discriminating motifs have a skewed positional distribution that exhibits extreme enrichment very close to the HMB boundaries.

**Figure 4.**
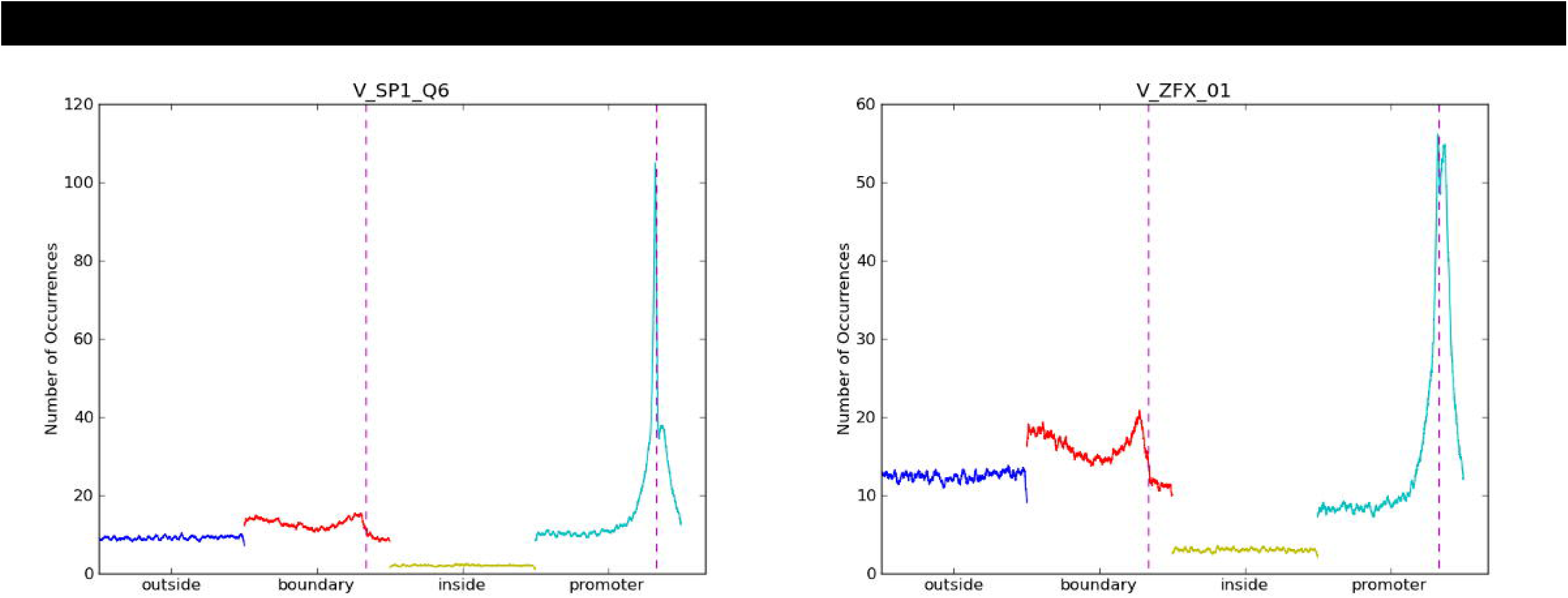
Positional profile of binding sites for ZFX and SP1. Number of occurrences in 100 bp windows as function of genomic distance to HMB or promoter start site for TFs ZFX_01 (a) and SP1_Q6 (b). The dotted vertical line indicates the location of HMB and promoter respectively. ‘Outside’ and ‘inside’ correspond to 6kb sized genomic regions outside or inside HMBs respectively.

### Characterization of the most discriminating Transcription Factor motifs

Several TFs are directly involved in histone modification and several more TFs are otherwise known to interact with chromatin modification enzymes [23]. We assessed whether the TFs whose motifs are most discriminative of HMB boundaries are involved in chromatin modification, either directly or by interacting with a chromatin modification enzyme. We first compiled a set of 492 genes annotated as Chromatin Modification Enzymes (CME) from the ENSMBL database. For each of the 931 TransFac motifs, we obtained the Ensemble Gene ID for the corresponding TF protein and then obtained the set of annotated proteins known to interact with the particular TF using the string-db R package, which is based on the STRING database of protein interactions [24], For each pair of regions compared (say, HMB boundary versus Promoter), we assessed whether the most discriminating motifs and their interacting partners are enriched for CMEs. To do so we obtained the top 20, 25, 40, and 50 motifs according to Mean Decrease Accuracy (see Methods), and compared the prevalence of CMEs among these motifs and their interacting partners against the rest of the available TF proteins as background. For each comparison, we assessed enrichment using Fisher’s Exact test. We found that the most discriminating TF motifs (Supplementary Table 1-3) in HMB boundaries and their interacting partners were enriched for CMEs relative to all other regions (inside HMB, outside HMB, and promoter regions, Table 2). Encouragingly, the fold enrichment of CMEs increases monotonically as we restrict ourselves towards more significant TFs, from top 50 to top 20 motifs only. These results suggest that relative to inside and outside regions, the HMB boundaries not only harbor distinct motifs but these motifs could also be responsible for distinct epigenetic profiles at HMB boundaries.

**Table 2.** Enrichment of chromatin modification enzymes among the most discriminating TF motifs and their interacting partners. Odds ratio (OR) and Fisher’s Exact test P-value for a chromatin modification enzyme enrichment test using the most discriminating (Top 20, 25, 40 or 50) TF binding site motifs for each classification task (as described in Table 1).

| Classification | Top 20 |  | Top 25 |  | Top 40 |  | Top 50 |  |
| --- | --- | --- | --- | --- | --- | --- | --- | --- |
|  | OR | P-Value | OR | P-Value | OR | P-Value | OR | P-Value |
| Boundary-Inside | 1.66 | 2.8e-9 | 1.57 | 6.1e-8 | 1.48 | 7.5e-7 | 1.46 | 7.8e-7 |
| Boundary-Outside | 1.61 | 3.0e-8 | 1.53 | 3.5e-7 | 1.50 | 2.4e-7 | 1.45 | 1.4e-6 |
| Boundary-Promoter | 1.64 | 7.6e-9 | 1.56 | 1.2e-7 | 1.44 | 3.9e-6 | 1.45 | 1.1e-6 |

Supplementary Table 5 lists the 135 CMEs that interact with the top 20 enriched motifs in each of the three comparisons – boundary versus inside, outside, and promoter. Interestingly, these 135 CMEs include two DNA methyltransferases DNMT3A/B, and also P300, which is a well-known marker of regulatory enhancers.

### Hypo-methylated blocks may be informed by chromatin structure

Our analysis so far suggests that the HMB boundary regions possess distinguishing genomic and epigenomic characteristics which may underlie their role as nucleation or termination of the methylation alteration. In addition, it is likely that the spread and confinement of epigenomic alteration within HMBs may be informed by preexisting chromatin organization and structure. This is suggested by a previous study that showed a significant overlap between cDMRs and LADs [5].

Based on Hi-C assay, which provides quantitative evidence of physical interactions between genomic loci, previous work has identified the so-called Topological Associating Domains (TAD), which are mega-base-sized genomic regions with a much greater interactions within the regions relative to across regions. TADs are relatively conserved across cell lines and species, and thus represent an underlying structural backbone of the chromatin. Based on 3,127 TADs reported in [16], we measured the proximity of each TAD boundaries to the closest HMB boundary, and compared the resulting positional distribution with that for a control set of randomly selected genomic loci. TAD boundaries are significantly closer (~43kb) in genomic distances to a HMB boundary compared with the expected ~71kb (ratio of mean = 3.8, ratio of median = 1.7, Wilcoxon test p-value = 5.4e-55, Figure. 5a).

**Figure 5.**
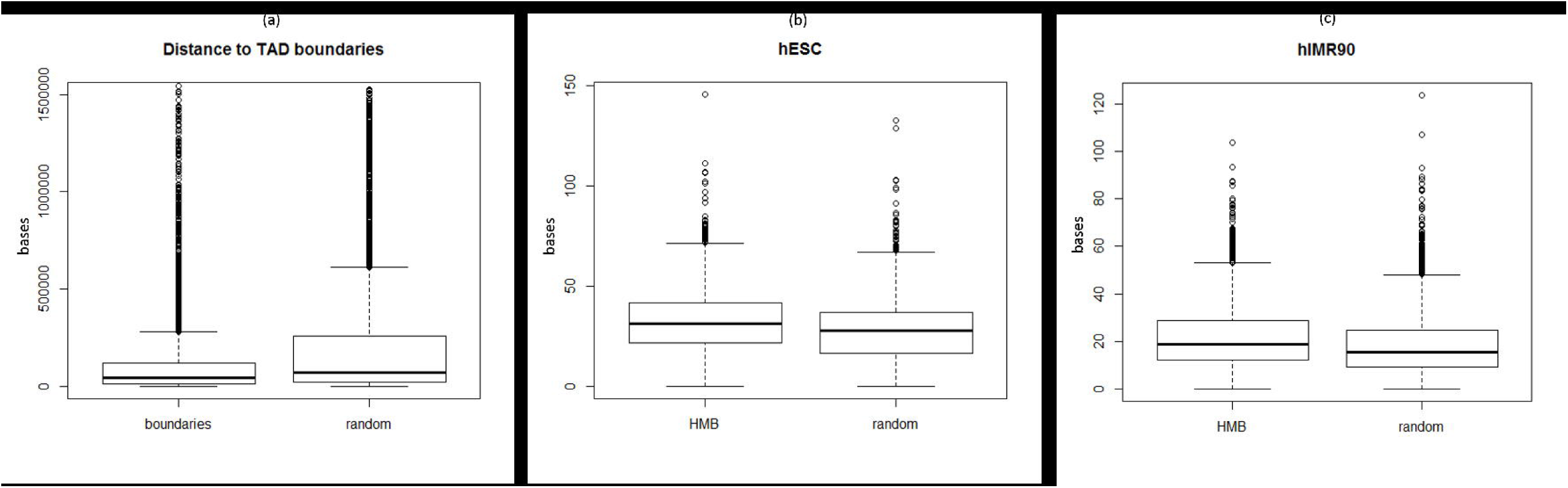
Hypo-methylated blocks associate with topological domains in the chromatin structure. ***(a)*** Boxplot of genomic distance (in bases) between TAD boundary (obtained from hESC) to nearest colon cancer HMB boundary. Distances between TADs and random 6kb genomic regions are included as background, **(b)** Boxplots of average Hi-C interaction for bins within HMBs in hESC, with average interactions randomly generated genomic regions of similar size and GC content included as background, **(c)** same as (b) and for IMR90 cell line.

Because TADs were identified based on a statistical overrepresentation of intra-region interaction, we also directly assessed using the Hi-C data, whether HMBs show an enriched intra-block interaction compared to inter-block interactions. Unfortunately Hi-C data is not available for human colon tissue. Based on the Hi-C data in hESC, and hIMR90 cell line (yuelab.org/hi-c/download.html), as shown in Figure 5b-c, we found a significantly greater interactions within HMBs compared to within random blocks controlled for length (For hESC: mean_HMB = 32, mean_Random = 27, Wilcoxon test p-value = 3.2e-38. For hIMR90: mean_HMB = 21.8, mean_Random = 18.3, Wilcoxon test p-value = 4.1e-34).

Overall, these analyses suggest that long domains of altered methylation in colon cancer may in part be informed by the underlying chromatin structure of the normal cell.

### CTCF binding sites coincide with the H3K4me3 signal in HMB boundaries

Among its numerous roles, CTCF is known to act as insulator by restricting the spread of heterochromatin, and is also involved in the maintenance of three dimensional chromatin conformation in part by stabilizing long-distance interactions [25]. Consistent with the role of insulator, CTCF binding sites are enriched between TADs [16]. We assessed whether CTCF binding sites are enriched near HMB boundaries. We downloaded the *in vivo* CTCF binding sites for colon cancer tissue from CTCFBSDB 2.0 database (insulatordb.uthsc.edu/). We found that HMBs were often bounded by CTCF binding sites. The frequency of CTCF in the 6 kb HMB boundaries (21%) was significantly higher than random blocks inside (14%) and outside (18%) HMBs, where the total number of regions was each set is ~20k. Moreover and interestingly, the HMB boundaries with a CTCF binding site had significantly higher levels of H3K4me3 signal than the boudaries without a CTCF binding site (ratio of mean = 1.4, Wilcoxon test p-value = 3.7e-24). Overall, this suggests that HMB boundaries are enriched for CTCF, as is expected for structural chromatin domains, but the presence of CTCF is in fact linked to the promoter-like characteristic of HMB boundaries.

## Discussion

In this study, we have characterized the regulatory landscape of large regions of methylation loss in colon cancer. We have found that the putative binding sites for specific TFs potentially involved in chromatin modification are distinguishing features of the DNA sequence at HMB boundaries. We also found that while activating histone marks common to promoters are enriched in HMB boundaries, HMB boundaries still show a distinct pattern of TF motif profile relative to known promoters. Finally, we found that the specific domains where HMBs occur are reflective of general chromatin organization of the normal cell.

Based on our qualitative assessment, we found that TFs enriched in HMB boundaries include those involved in demethylation, cell proliferation and cell cycle, hallmarks of cancer. For instance, for the most discriminative motif Sp1, high expression of Sp1 is known to disrupt cell cycle. Sp1 deregulation might be beneficial for tumor cells and its overexpression is known to induce apoptosis of untransformed cells [26]. Other members of Sp TF family also play roles in metastasis and growth of different tumor types [27]. In our analysis, multiple TFs from this family were found to be enriched in HMB boundaries. Zfx presents another illustrative example, as it controls the self-renewal of embryonic and adult hematopoietic stem cells [28]. Zfx also controls BCR-induced proliferation and survival of B lymphocytes [29]. Another detected TF FoxO is central to the integration of growth factor signaling, oxidative stress and inflammation, and is involved in tumor suppression [30] and DNA demethylation process in B-cell development [31]. Finally, TF Zfp281 is known to play a role in cell pluripotency [32], chromatin remodeling [33], and inhibition of nanog auto-repression [34],

Loss of methylation in large domains has been identified as a consistent and stable mark in solid tumors [5, 35]. While the degree of methylation loss increases with tumor progression, intra-sample variability in DNA methylation and gene expression is greater within these domains [35]. These findings point to a general loss of epigenomic and transcriptomic stability that is essential to the normal behavior of the cell. The co-localization of these domains with lamin-associated domains [5], with TADs (as found in this study), and the enrichment of CTCF binding in the boundaries of these domains suggest that a loss of chromatin organization is concomitant with this loss of epigenomic and transcriptomic stability.

We note a few limitations of our analyses. Our analyses are based on 6 kb region flanking the HMB boundary. This choice, while reasoned, is somewhat arbitrary. Although our analyses suggest that HMB formation is associated with specific genomic, epigenomic, and chromatin features, it does not clarify the causality leading from TF binding to hypomethylation and ultimately to the previous observed aberrant gene expression in HMBs. While we observed specific patterns of certain epigenomic marks at HMB boundaries, these may be ultimately a reflection of the genomic characteristics [36]. Moreover our analysis is based on putative binding site and not based on *in vivo* binding data for the TFs, which are currently not available for a majority of TF. Nevertheless, our analyses do suggest a potential link between specific genomic marks and HMB boundaries, which require future experimental studies of the underlying mechanisms.

## Conclusion

Taken together, our analyses suggest that the overall architecture of HMBs is guided by pre-existing chromatin architecture, while their creation in cancers may be caused by aberrant activity of promoter-like sequences at the boundary. Our results are consistent with a model where a loss of chromatin organization and a concomitant loss of epigenetic stability make previously inaccessible TF binding sites accessible for proteins involved in chromatin modification as well as cellular fate, whose binding sites are enriched within domains of inaccessible chromatin where HMBs reside. The binding of specific DNA binding factors at HMB boundaries may further participate in methylation loss.

## Authors’ contributions

HCB and SH conceived and designed the project. M.S. performed all the analyses. All authors helped write the manuscript.

## Acknowledgements

This work was supported by NIH R01GM100335 to S.H.

## Additional Information

**1. Additional File 1 contains supplementary figures 1-3.**

**Supplementary Figure 1.** *Pattern of histone marks near HMB boundaries: (a) H3K4me1, (b) H3K9me3, (c) H3K27me3, (d) H3K36me3.*

**Supplementary Figure 2.** ROC for HMB boundaries versus inside/outside for the test set.

**Supplementary Figure 3.** *Frequency plots for the TF motifs listed in Supplementary Table 4.*

**2. Supplementary Tables1-5.** Legends are provided along with the tables.

